# Large-scale dendritic spine extraction and analysis through petascale computing

**DOI:** 10.1101/2021.07.29.454371

**Authors:** Gregg Wildenberg, Hanyu Li, Griffin Badalamente, Thomas D. Uram, Nicola J. Ferrier, Narayanan Kasthuri

**Affiliations:** University of Chicago, Department of Neurobiology, Chicago, IL USA; Argonne Leadership Computing Facility, Argonne National Laboratory, Lemont, IL USA; Mathematics and Computer Science Division, Argonne National Laboratory, Lemont, IL USA

**Keywords:** connectomics, electron microscopy, automatic segmentation, synapses, mouse brain, machine learning

## Abstract

The synapse is a central player in the nervous system serving as the key structure that permits the relay of electrical and chemical signals from one neuron to another. The anatomy of the synapse contains important information about the signals and the strength of signal it transmits. Because of their small size, however, electron microscopy (EM) is the only method capable of directly visualizing synapse morphology and remains the gold standard for studying synapse morphology. Historically, EM has been limited to small fields of view and often only in 2D, but recent advances in automated serial EM (“connectomics”) have enabled collecting large EM volumes that capture significant fractions of neurons and the different classes of synapses they receive (i.e. shaft, spine, soma, axon). However, even with recent advances in automatic segmentation methods, extracting neuronal and synaptic profiles from these connectomics datasets are difficult to scale over large EM volumes. Without methods that speed up automatic segmentation over large volumes, the full potential of utilizing these new EM methods to advance studies related to synapse morphologies will never be fully realized. To solve this problem, we describe our work to leverage Argonne leadership-scale supercomputers for segmentation of a 0.6 terabyte dataset using state of the art machine learning-based segmentation methods on a significant fraction of the 11.69 petaFLOPs supercomputer Theta at Argonne National Laboratory. We describe an iterative pipeline that couples human and machine feedback to produce accurate segmentation results in time frames that will make connectomics a more routine method for exploring how synapse biology changes across a number of biological conditions. Finally, we demonstrate how dendritic spines can be algorithmically extracted from the segmentation dataset for analysis of spine morphologies. Advancing this effort at large compute scale is expected to yield benefits in turnaround time for segmentation of individual datasets, accelerating the path to biology results and providing population-level insight into how thousands of synapses originate from different neurons; we expect to also reap benefits in terms of greater accuracy from the more compute-intensive algorithms these systems enable.

## 1 INTRODUCTION

Synapse morphology conveys a wealth of information about its function. At the most rudimentary level, whether the synapse is located at a dendritic spine or shaft of an excitatory neuron identifies whether that synapse transmits excitatory or inhibitory signals, respectively (Piergiorgio Strata, 1999). Additionally, the size of the dendritic spine head and area of the post-synaptic density (PSD) is linearly correlated with the strength of the synapse (S et al., 2021). Lastly, at even finer detail, the presence or absence of a spine apparatus or mitochondria, and size of neurotransmitter vesicles also provides indirect information about the strength and type of synapse including neurotransmitter type and long term potentiation (LTP) (Segal, 2010; P et al., 2008; Bourne JN, 2008; Edwards, 1998).

Despite its importance and ability to draw correlations between synapse structure and function, studying synapse morphology at scale (i.e. analyzing the morphology of hundreds to thousands of synapses) has been difficult because electron microscopy is required to visualize synapses directly and, until recently, many EM techniques were restricted to single 2D sections or a small series of sections centered on a small field of view (FOV) where only a handful of synapses are imaged. These small FOV images often leave out critical information including the location of the synapse along the postsynaptic target (e.g. apical vs distal dendrite) and the identity of the postsynaptic target (e.g. Layer 2/3 versus Layer 4 pyramidal neuron). Moreover, the inability to image and classify synapses at scale across numerous neurons has severely limited our appreciation of how different morphological features of synapses vary across a population of synapses.

Most notably, a recent analysis between synapse structure and function discovered a linear correlation between the amplitude of the excitatory postsynaptic potential (EPSP) (i.e. strength of synapse activity) and the the area of the postsynaptic density (PSD) and dendrite spine volume (Holler, 2021). Similarly, visual deprivation results in lower variation in axonal bouton size suggesting that similar morphological changes occur on the presynaptic side of synapses as well (Sammons et al., 2018). These add to the accumulating evidence that changes in activity correspond with changes in anatomy creating the possibilty of mapping synaptic strength directly onto synapse morphology. While these and other studies (Araya et al., 2014, 2006; Borczyk et al., 2019) establish important structure/function relationships in synapse anatomy, these analyses are often limited to a small number of synapses on a single neuron, leaving many open questions on how these anatomical measurements scale different dendritic locations (i.e. distal vs proximal dendrites), at different regions of the neuron (dendrite spine and shaft, soma, or axon), and across populations of neurons (Harris and Weinberg, 2012).

Automated serial EM (“connectomics”) has rapidly emerged as a critical tool for capturing a large number of synapses across numerous neurons (Kasthuri, 2015; Hayworth et al., 2020; Januszewski et al., 2018; Bae et al., 2018; Baena et al., 2019; Motta et al., 2019a; Yin et al., 2020). The high resolution (∼3-6nm), large fields of view capable of being captured over a 3D volume makes connectomic approaches the ideal solution for studying synapse morphology at scale. However, methods to identify different biological features primarily rely on laborious, human-intensive tracings to segment objects within the EM volumes. Thus, while imaging larger volumes has advanced rapidly, there are significant bottlenecks at the level of segmentation that have hampered progress.

Recent advances in automatic segmentation have greatly aided in the process of extracting neuronal features from connectomic datasets. However, these algorithms are presently very slow and require significant computational resources to the point where they are impractical for many research groups. The Department of Energy (DOE) National Laboratories have historically provided groundbreaking infrastructure to academic research groups. In 1995, Argonne National Laboratory constructed the Advanced Photon Source (APS) used for high resolution x-ray imaging for research in numerous fields including from material sciences, biology, physics, and chemistry. In 1997, the DOE created the Joint Genome Institute (JGI) to unite expertise and technology to facilitate the rapid advancement of whole genome sequencing and since 2004, the JGI has served as a user facility for genomics research. Central to many national labs is the construction and operation of advanced supercomputers to facilitate user-based large scale simulation and analysis of data generated both within and outside the national labs. Continuing this trend, Argonne National Laboratory will soon deploy an exascale supercomputer, capable of performing one billion billion floating point operations per second. Thus, in addition to having the most powerful computers in the country, the DOE national laboratories have unparalleled infrastructure for processing, storing and moving large data.

We describe our effort to leverage the supercomputers at the Argonne Leadership Computing Facility (ALCF) to bridge the gap between the field of connectomics and the large scale computing necessary for fast automatic segmentation of EM volumetric data sets. Specifically, we show how our parallelization of existing segmentation algorithms increase segmentation throughput many-fold, and present a workflow for reconstructing volumes of dendrite spines for spine volume analysis with custom, open-source scripts. Finally, we discuss the future goals of developing this pipeline as a user-based platform for high performance computing of future connectomics datasets.

## 2 MATERIALS AND METHODS

### 2.1 Tissue preparation and EM imaging

A mouse brain was prepared in the same manner and as previously described (Hua et al., 2015). Briefly, an anesthetized animal was first transcardially perfused with 10ml 0.1 M Sodium Cacodylate buffer, pH 7.4 (Electron Microscopy Sciences [EMS]) followed by 20 ml of fixative containing 2% paraformaldehyde (EMS), 2.5% glutaraldehyde (EMS) in 0.1 M Sodium Cacodylate buffer, pH 7.4 (EMS). The brain was removed and placed in fixative for at least 24 hours at 4°C. A series of 300 µm vibratome sections were prepared and put into fixative for 24 hours at 4°C. The primary visual cortex (V1) was identified using areal landmarks and reference atlases. A small piece (∼2 × 2 mm) containing V1 was cut out and prepared for EM by staining sequentially with 2% osmium tetroxide (EMS) in Sodium Cacodylate buffer, 2.5% potassium ferrocyanide (Sigma-Aldrich), thiocarbohydrazide, unbuffered 2% osmium tetroxide, 1% uranyl acetate, and 0.66% Aspartic acid buffered Lead (II) Nitrate with extensive rinses between each step with the exception of potassium ferrocyanide. The tissue was then dehydrated in ethanol and propylene oxide and infiltrated with 812 Epon resin (EMS, Mixture: 49% Embed 812, 28% DDSA, 21% NMA, and 2.0% DMP 30). The resin-infiltrated tissue was cured at 60°C for 3 days. Using a commercial ultramicrotome (Powertome, RMC), the cured block was trimmed to a 1.0mm x 1.5 mm rectangle and 2,000, 40nm thick sections were collected on polyimide tape (Kapton) using an automated tape collecting device (ATUM, RMC) and assembled on silicon wafers as previously described (Kasthuri, 2015). A ROI of about 100 µm^2^ surrounding Layer 4 was selected for high resolution imaging and 913 of the 2,000 sections were imaged. Sections were acquired using backscattered electron detection with a Gemini 300 scanning electron microscope (Carl Zeiss), equipped with ATLAS software for automated wafer imaging. Dwell times for all datasets were 1.0 microsecond.

### 2.2 Computing Resources

This work leveraged computing resources at Argonne National Laboratory: Theta, an 11.69 PFLOPs supercomputer, and Cooley, both at the Argonne Leadership Computing Facility. Theta is composed of 4,392 CPU-based compute nodes, each with a 64-core, 1.3-GHz Intel Xeon Phi 7230 processor, 192GB DDR4 RAM and 16GB high-bandwidth MCDRAM, and 24 GPU-based compute nodes, each with two 24-core AMD Rome CPUs, eight NVIDIA A100 GPUs, and 1 TB RAM. The Cooley analysis cluster consists of 126 compute nodes, each with two 6-core Intel E5-2620 CPUs, one NVIDIA K80 (with two GK210 GPUs), and 384GB RAM.

### 2.3 Automated segmentation

Segmentation of neuronal objects for analysis consists of identifying individual objects by recognizing boundaries to disambiguate them from neighboring objects, to collect pixels belonging to each object, and uniquely labeling objects for later analysis. For segmentation of objects in electron microscopy image stacks, we used the Flood Filling Network (FFN) (Januszewski et al., 2018) developed by Google. FFN is a neural network trained on user annotations that identify boundaries in a sample of the input data. The network consists of a series of blocks, with each block comprising a three-dimensional convolutional layer, a rectified linear activation layer (ReLU), a second convolutional layer, and a second ReLU layer; the input to the convolutional block is also connected to the final ReLU layer by a skip connection. Convolutional layers use the SAME mode, and no pooling operations are used in the network, thereby preserving the input size. During training, FFN predicts the probability that pixels within a subvolume belong to an object, for a collection of subvolumes at predetermined seed points. As the network trains, it moves adaptively through the volume by translating the FOV by a prescribed delta in [x,y,z]. With each predicted FOV, the network weights are updated by stochastic gradient descent, using a voxelwise log-loss computed against the input training data. Once trained to an acceptable accuracy, measured using the F1 score, the trained model is run in a forward pass over the entire volume to produce a full segmentation. In the case of distributed inference over a set of compute nodes, FFN will produce a set of individually segmented subvolumes, which are reconciled to produce a consistently segmented full volume.

### 2.4 Dendritic Spine separation and analysis

Isolation of individual segmented spiny dendrites and subsequent separation of spines from the dendrite was performed using homemade scripts using open source Python libraries. Spine separation was performed with computer vision techniques relying on the thinness of the dendritic spines compared to the width of the dendrite and the bouton. Eroding the full structure severed the thin spines from the dendrite, allowing for isolated identification of the dendrite. Dilating the dendrite structure back to its original size and removing it from the original dataset left only the spines and boutons. These spines were then uniquely labeled and analyzed as individual objects. The analysis done here is simply comparison of spine volumes (done by converting voxel volumes into micrometer volumes) but more varied morphological analysis is possible. All code was assembled into Jupyter Notebooks that are available upon request.

## 3 RESULTS

We prepared 300µm thick sections of ‘adult’ mouse primary visual cortex (V1) (105 days old, male) for nano-scale EM-based connectomics. V1 was identified using areal and cellular landmarks and the Allen Brain Institute online reference atlas. Tissue samples were prepared as previously described (Hua et al., 2015) (and see Methods). We first collected 3,000 ultra-thin sections using the ATUM approach (Kasthuri, 2015) from mouse (each section: 0.8mm x 1.5 mm x 40nm) where each individual section spanned all cortical layers. We identified Layer 4 (L4) using previously described landmarks (Motta et al., 2019b) and imaged a volume of 100 µm x 100 µm x 36 µm at 6nm in-plane (xy) resolution for synaptic -level automated segmentation and analysis. Tiles comprising individual serial sections were then stitched in 2D and the stitched images were 3D-aligned using TrakEM2 (Cardona et al., 2012) and AlignTK (https://mmbios.pitt.edu/software) which were both run in parallel for fast alignment, ultimately resulting in a fully aligned EM stack in approximately one week. Figure 1A shows an example of a stereotypical spine synapse identified in our EM dataset hallmarked by the presence of a postsynaptic density (PSD) and numerous vesicles in the presynaptic axon. Because our dataset is volumetric, the full 3D architecture of the synapse is able to be manually reconstructed (Figure 1B).

**Figure 1.**
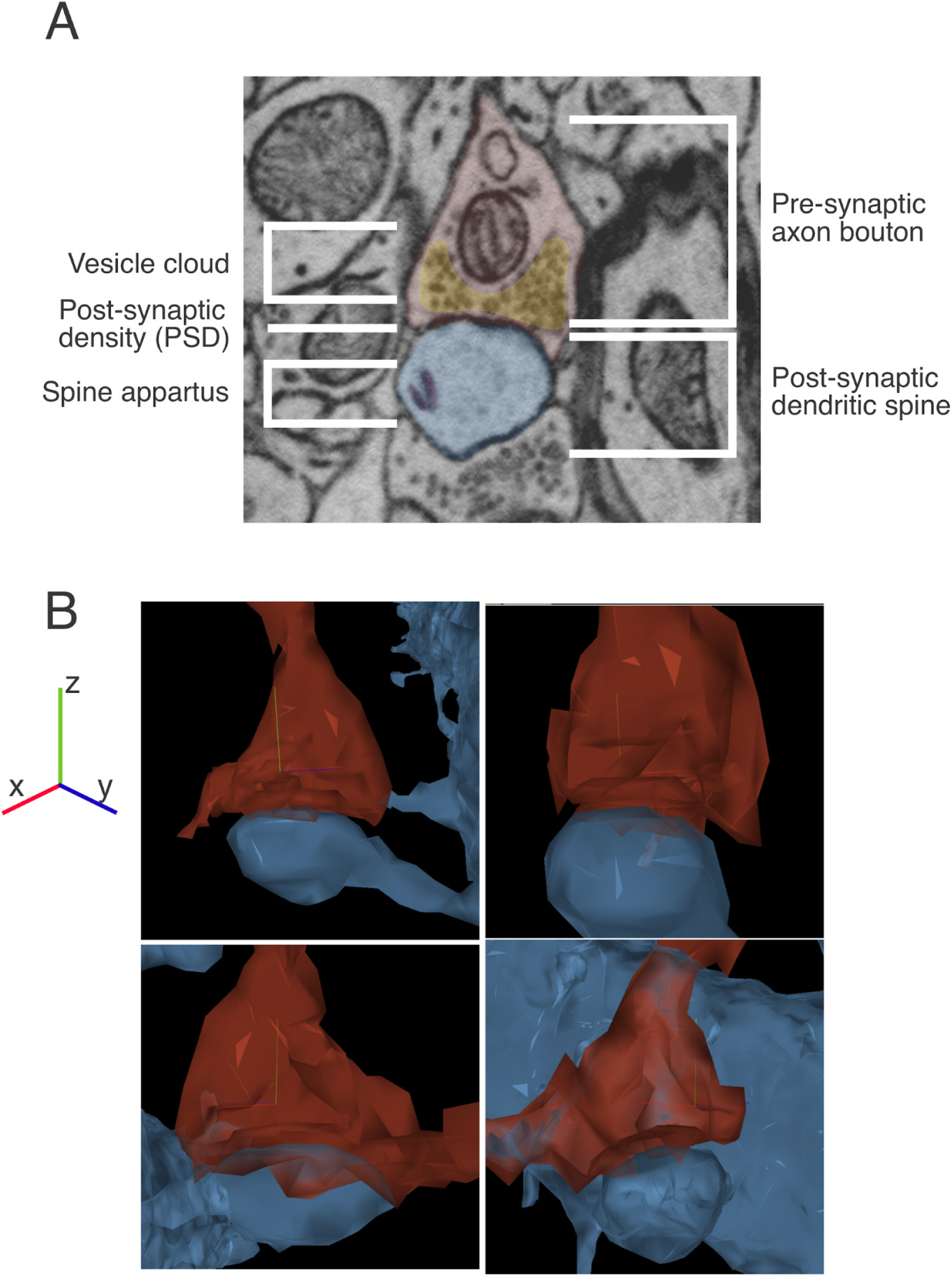
Core structure of the synapse. **(A)** EM image of an excitatory synapse onto a dendritic spine. The presynaptic axonal bouton (red), hallmarked by the presence of a vesicle cloud (yellow), makes physical contact with the postsynaptic dendritic spine (blue) through the establishment of an electron dense postsynaptic density (PSD). In some cases, the postsynaptic dendritic spine also contains a membrane structure called the spine apparatus. **(B)** Different angular views showing a 3D reconstruction of a dendritic spine (blue) receiving a synapse (red) from EM images shown in (A).

Because the data captures a large portion of neurons (i.e. soma, dendrites, axons), different classes of synapses can be readily identified based on where they are located along the post-synaptic target providing information about whether they are likely excitatory or inhibitory synapses (Piergiorgio Strata, 1999). Additionally, for inhibitory synapses on the shaft and soma, the location of the synapse predicts the most likely cell type of the presynaptic neuron (e.g. Chandelier cells exclusively make axo-axonic synapses) (Wang et al., 2016; Kepecs and Fishell, 2014) (Figure 2). Annotating a small number of synapses and post-synaptic targets in connectomic datasets is manageable, however, to scale annotation where thousands of synapses can be studied across hundreds of neurons is prohibitively slow. Recent developments in machine learning automatic segmentation (Januszewski et al., 2018; Funke et al., 2019; Lee et al., 2019) have made it possible to automate the laborious and slow process of manual segmentation; implementing these algorithms at scale where terabyte-scale datasets can be rapidly segmented remains a major hurdle to fast processing and analysis of connectomic data.

**Figure 2.**
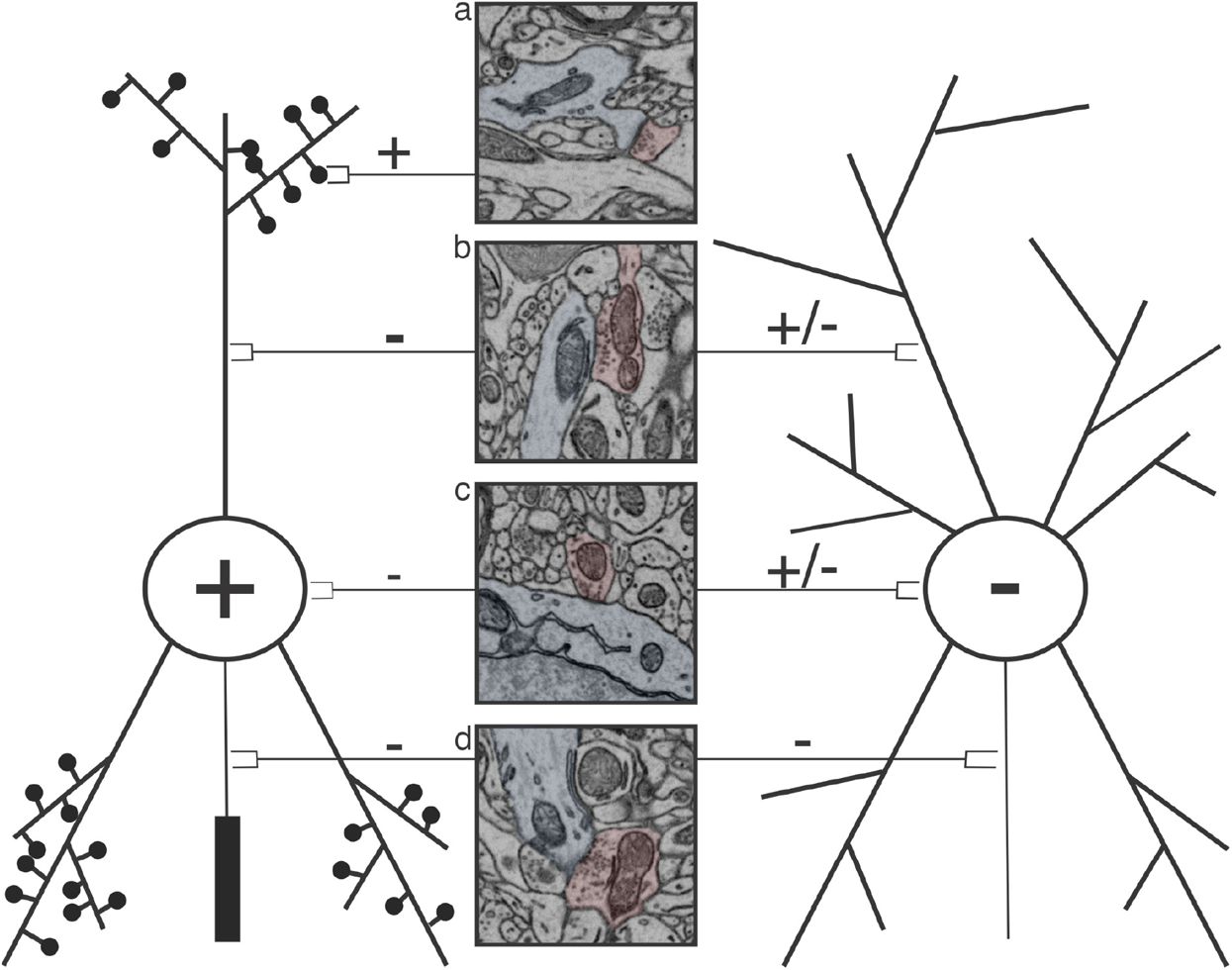
Schematic showing a stereotypical excitatory (+) (left) and inhibitory (-) neuron (right) with corresponding 2D EM images showing the postsynaptic target area (blue) of different synapses (red) each kind of neuron receives. Excitatory neurons receive excitatory synapses on dendritic spines (a) and inhibitory synapses on their dendritic shaft (b), soma (c), and axon (d). Inhibitory neurons receive excitatory or inhibitory synapses on their shaft (b) or soma (c) and inhibitory synapses on their axon (d).

We sought to overcome this hurdle by using Argonne National Laboratory supercomputers to construct an algorithmic pipeline for automatic segmentation to determine whether national laboratory supercomputers are a suitable platform for ingesting published algorithms and increasing the rate at which connectomic datasets can be segmented. We chose the flood filling network (FFN) (Januszewski et al., 2018) segmentation code which has been implemented in multiple, large volume connectomic datasets derived from different species and different connectomics technologies (i.e. ATUM and SEM vs manual section pickup and TEM) (Shapson-Coe et al., 2021; Li et al., 2020) demonstrating that FFN is applicable to a wide range of connectomic studies and thus ideally suited to serve a broad range of samples.

In the current work, we used an FFN network configured with nine convolutional blocks (eighteen convolutional layers), FOV size 33 × 33 × 17, and FOV delta 8 × 8 × 4. In this configuration, the network had 0.5 million trainable parameters. Acknowledging the human-intensive process of annotating objects to establish training data sets, we started with an FFN model that was pre-trained on data from Kasthuri Kasthuri (2015) (K11) to attempt to reduce the human time involved in training the model from scratch. After an initial evaluation of the segmentation produced by inferencing with the K11 model on the current data, we completed several more rounds of annotating data using WebKnossos Boergens et al. (2017), incrementally retraining the FFN model and evaluating segmentation performance on test data until an acceptable accuracy was achieved, at which point we could proceed to large-scale inference using this model. This incremental approach of annotation, training, and inference required approximately thirty person-hours for annotation within a 512 × 512 × 128 pixel subvolume.

Segmentation involves leveraging the trained model for inference over the entire input volume. To conduct inference in parallel over the large volume, we decompose the volume into subvolumes to be segmented by individual compute nodes, and subsequently reconcile the collection of subvolumes into a single monolithic volume with consistently labeled objects. To this end, it is important to consider the performance of the model at this scale. To optimize performance of inference with FFN on the Intel KNL processors, we conducted a scaling study of FFN on a single compute node of the Theta supercomputer. For this study, we began with a subvolume of size 688 × 163 × 50, and ran FFN inference on a single Theta compute node with an increasing number of threads. Details from this study are shown in Table1, where it is apparent that FFN can fully utilize the 64 processor cores in the KNL processor, segmenting the full subvolume in 15 minutes. Notably, while each KNL core is equipped with four hardware threads, performance unfortunately degrades when using two hardware threads per core (128 threads total) and four hardware threads per core (256 threads).

**Table 1.** FFN Inference time on KNL 7230 processor with varying numbers of threads, with a subvolume of size 688 × 163 × 50 voxels. The best performance is obtained with 64 threads (one thread per core); utilizing the multiple hardware threads per core beyond this point delivers no advantage.

| Number of Threads | FFN Inference time (minutes) |
| --- | --- |
| 4 | 32 |
| 8 | 30 |
| 16 | 24 |
| 32 | 20 |
| **64** | **15** |
| 128 | 23 |
| 256 | 42 |

The volume used for the current study consisted of a subset of the overall data, 913 sections, with each section of extent 21020 × 23237 pixels. Inference/segmentation was run in two computing scenarios to compare performance on the Theta KNL and Theta GPU nodes; we report simple statistics of these runs here, while reserving a more detailed, larger-scale study for future publication. In the first scenario, on Theta KNL nodes, each inference task was assigned a subvolume of 512 × 512 × 128 voxels, and was assigned 64 cores. The largest inference task was performed in an 882-node job that ran for 70 minutes and completed 1,142 subvolumes. This was followed by a series of smaller jobs to accommodate variability in runtimes and to optimize throughput in light of queue wait times on Theta. The average per-node voxel segmentation throughput for the entire volume in this scenario was 30 million voxels per node-hour.

In the second scenario, using Theta GPU nodes, each inference task was run on a single NVIDIA A100 GPU, with a subvolume of 512 × 512 × 256 voxels, for a total of 2990 subvolumes. The largest inference task in this scenario used 96 A100 GPUs for nine hours, segmenting 1070 subvolumes. The average inference runtime for these subvolumes was 49 minutes, leading to a per-GPU voxel segmentation throughput of 82 million voxels per GPU-hour. This GPU configuration was used to segment the full stack of 913 sections as reflected in the figures.

After completing full segmentation of the EM volume, the CloudVolume-formatted data can be manually inspected in Neuroglancer^1^ and manually parsed into individual objects depending on the desired analysis. To illustrate the utility of saturated segmentation for synapse analysis, we outline an example of how a user can go from segmentation to analysis of dendritic spine volume, but similar analyses can be seamlessly integrated from the point of a segmented volume rendered on Neuroglancer. We chose spine volume because the size of the dendritic spine has been correlated with synaptic strength (Borczyk et al., 2019) and we thus sought to determine the variability in dendritic spine volume as a coarse proxy to synaptic strength variability. First, because the fully segmented data is loaded into neuroglancer, it can be easily shared to multiple users enabling collaborations across multiple universities. In Neuroglancer, the full field of view can be visualized at a reduced resolution (Figure 3A, top left image), and once a target region is identified (white box), the user can zoom in on the data at different magnifications to see further detail (Figure 3A, m1, m2, m3). Once desired objects are identified, double clicking on individual objects brings up the full 3d rendering of that object. Figure 3B shows the combined renderings of all objects contained within the full resolution FOV shown in Figure 3A, m3.

**Figure 3.**
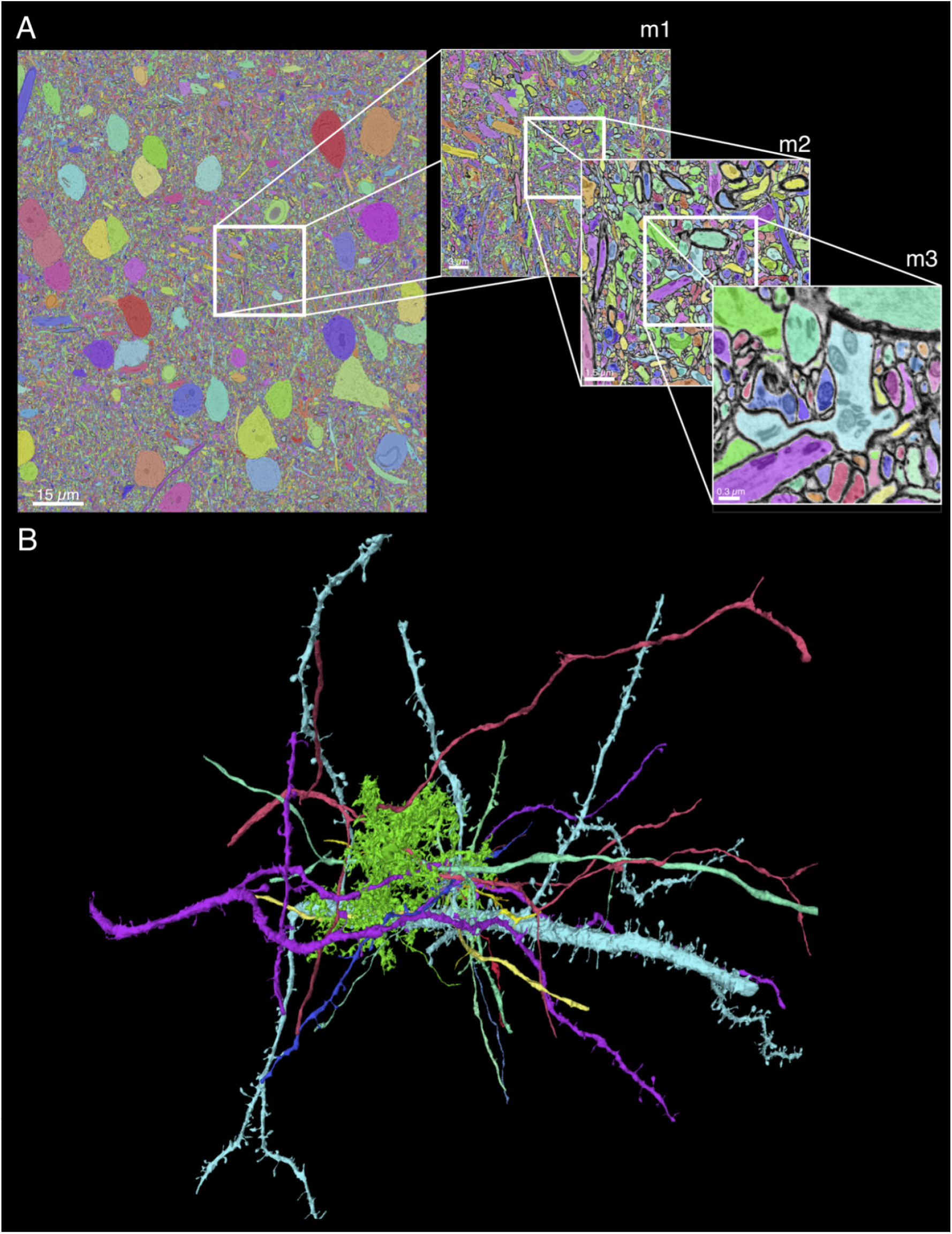
Output of scalable automatic segmentation of connectomic EM dataset. **(A)** Left: A single EM section with segmentation overlaid. Each color corresponds to a unique numerical ID assigned to each neuron. White boxes and corresponding images highlight a region cropped out to illustrate how using Neuroglancer, the data can be traversed to increasing magnifications (m1, m2, m3). **(B)** 3D rendering of objects contained within final field of view in (A, m3).

Once neurons of interest are identified, we then developed a script to extract those objects as a TIFF stack or HDF5 file from the segmentation using Python. Figure 5 illustrates three classes of postsynaptic targets extracted from the full volume. In the top panel, we identified the spines of three spiny dendrites. Spines can be easily identified in the raw EM dataset by looking for typical signatures of dendritic spines, including a spine apparatus, and following the spine through the volume to its connection back to the dendrite. Though not a focus of our current analysis pipeline, a similar process can be applied to smooth dendrites (presumably inhibitory neurons) and soma (Figure 4). However, because smooth dendrites and soma do not have spines, extracting the presynaptic bouton poses a different computational challenge that we will address in future iterations of our automated analysis methods. After extracting individual spiny dendrites from the segmentation volume, we ran these dendrites through a computer vision inspired algorithm we developed that separates the spines from the dendrite and assigns each spine a unique ID. This algorithm relies on erosions and dilations, which are two of the fundamental operations in morphological image processing. Erosion essentially shrinks the selected object in each dimension from the exterior. This removes the thinnest parts of a spine, separating boutons from dendrite. Dilation is not an inverse of erosions, but can be thought of as such. A dilation adds a layer on to the exterior of a selected object. While a dilation does not recover many of the fine details lost during an erosion, applying dilations to an eroded object can recover the general shape and size. As unique IDs, each spine can be mathematically analyzed as separate objects whereas in the prior step (i.e. before separation from the dendrite) they were regarded as a single object. We first demonstrate this pipeline on a single extracted dendrite (Figure 6A-D) then show how it works for an arbitrary number of dendrites (Figure 6E-F). The pipeline requires the TIFF stack or an HDF5 file containing neurons of interest extracted using the scripting described above as well as unique IDs for each desired dendrite. First, the pipeline reads the full volume into an array, removes everything that does not match the IDs given, and then converts the volume into a sparse representation of the data using the Sparse library (Abbasi, 2018). Figure 6A, left panel shows an example of a single dendrite extracted from the segmentation dataset (i.e. Figure 3). This sparse representation is saved to disk, allowing the second part of the pipeline to run multiple times without needing to read the full volume again, saving time and memory. The second part of the pipeline erodes the data a number of times (the data presented here is based on 10 erosions). Erosion is implemented with scikit-image (van der Walt et al., 2014). After erosions, the n largest objects are selected (where n is the number of selected dendrites). These n objects are virtually always the large dendrites, now missing the spines, which have been eroded away and no longer attached to the dendrite. These dendrites are then dilated (also implemented with scikit-image) back to their original size. Generally, this requires double the number of erosions that were applied (the data presented here is based on 20 dilations). These dendrites are then subtracted from the original data, leaving only the spines that were cut off from the full dendrite in the erosion process. Then, a simple size threshold was applied such that anything larger than 5.04e+8 nm^3^ or smaller than 8.64e+6 nm^3^ is removed from the data (Figure 6A, right panel). We used thresholds from previous reports (Arellano et al., 2007), but adjusted to give more range relative to the maximum and minimum spine volumes reported. It is important to note that dilation and erosion are not perfect inverses, so inconsistencies do occur. When we manually inspected the results to estimate the number of objects erroneously separated as spines (i.e. small protrusions in the dendrite that were algorithmically eroded off the dendrite, “False positives”), spines that did not get separated (i.e. “False negatives”), and spines successfully separated (i.e. “True”), we find our method has a combined false positive and negative error rate of 17% and was able to capture 83% of the total spines (Figure 6B). Once spines are separated, morphological analyses can be performed such as evaluating the distribution of spine volumes (Figure 6C). We then further modified our code so that an arbitrary number of dendrites can be simultaneously run to separate their spines into unique object IDs (Figure 6D).

**Figure 4.**
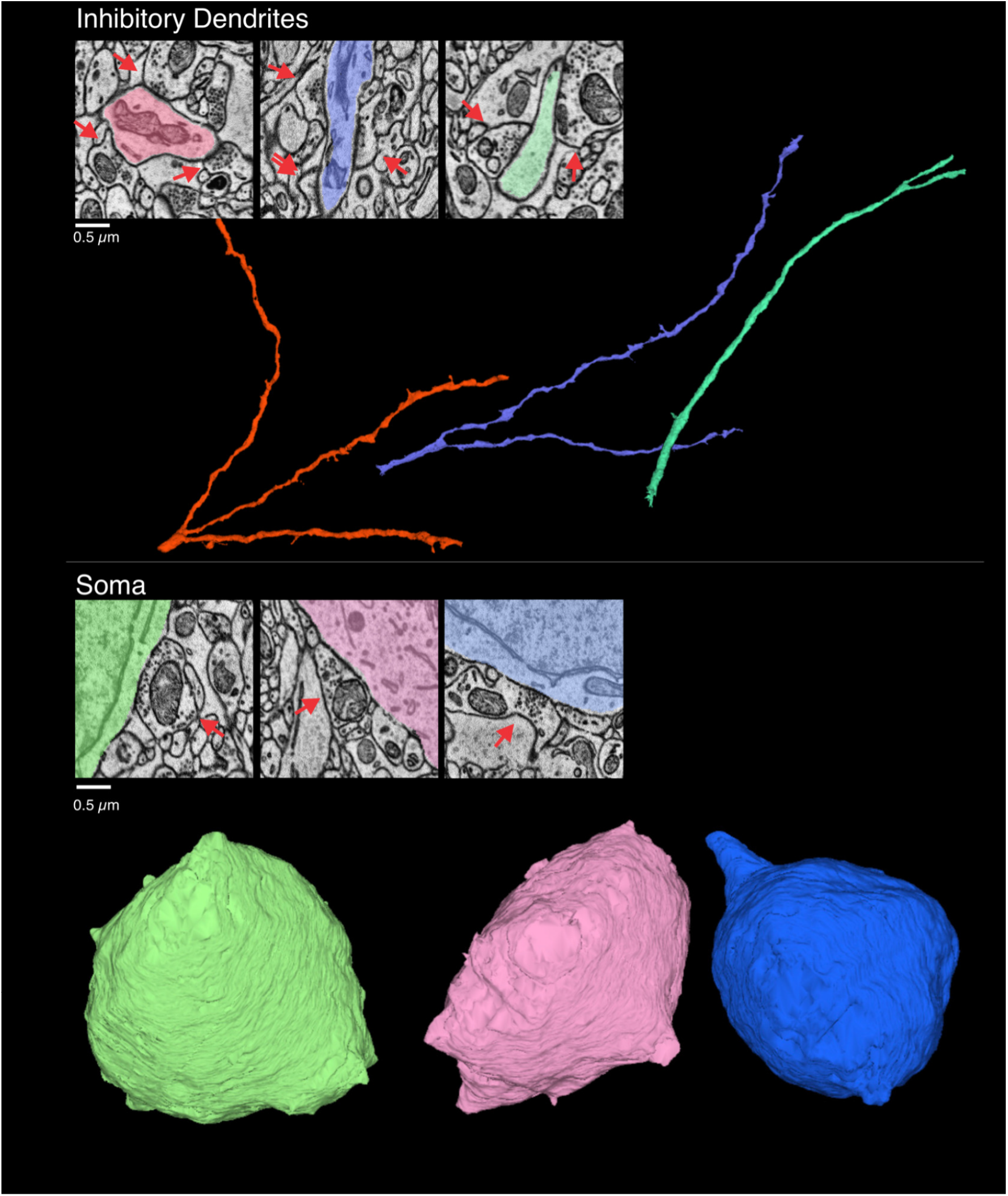
Individual aspinous dendrites and soma fragments isolated from dense segmentation. Top panel: 2D EM of three segmented aspinous dendritic shafts and corresponding full dendrite rendering. Bottom panel: 2D EM of three segmented soma and corresponding full rendering.

**Figure 5.**
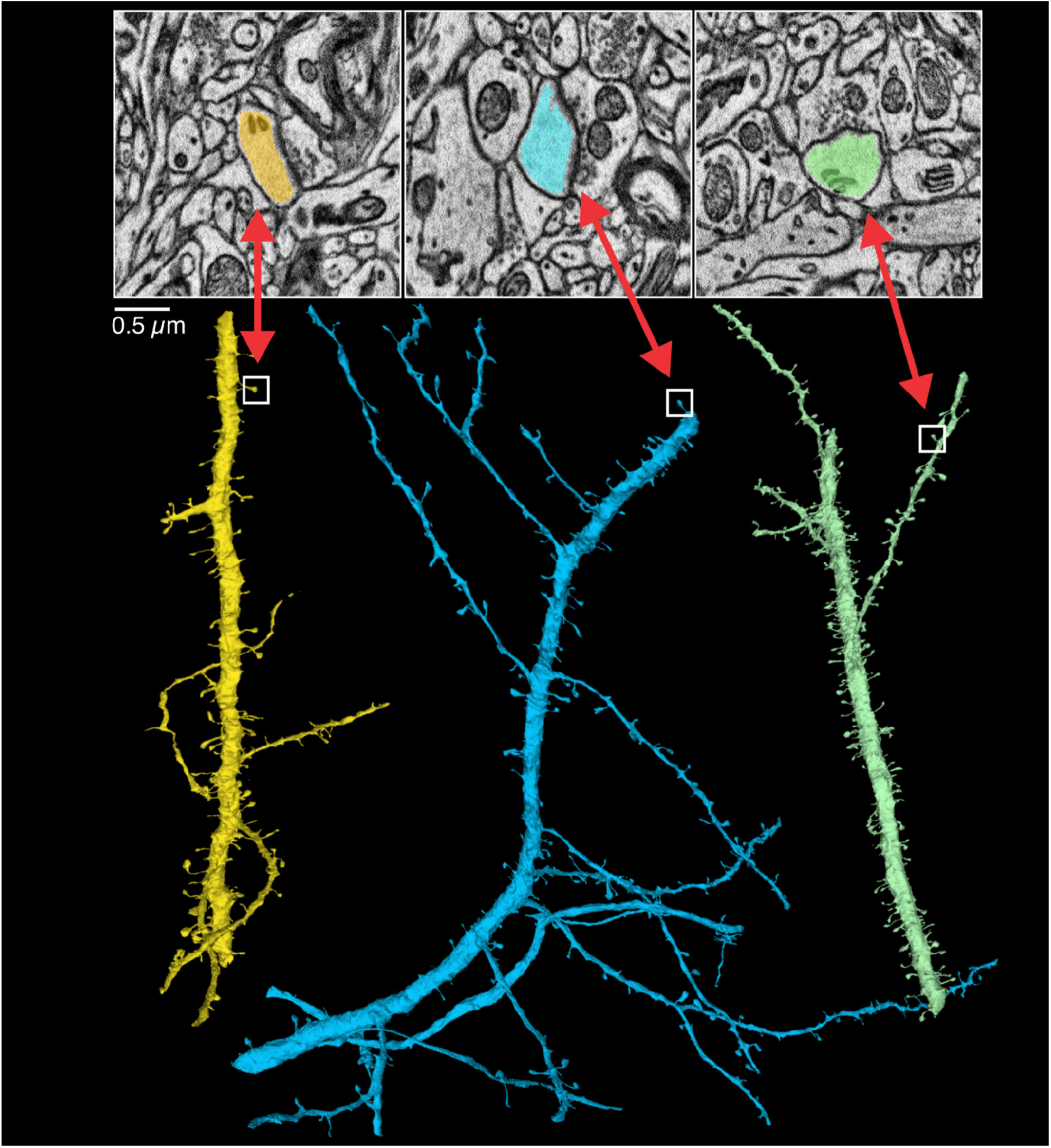
Individual spiny dendrites isolated from dense segmentation. 2D EM images of 3 segmented dendritic spines and corresponding full dendrite rendering the spines come from. Red arrows point to the postsynaptic spine and corresponding spine in full dendrite rendering.

**Figure 6.**
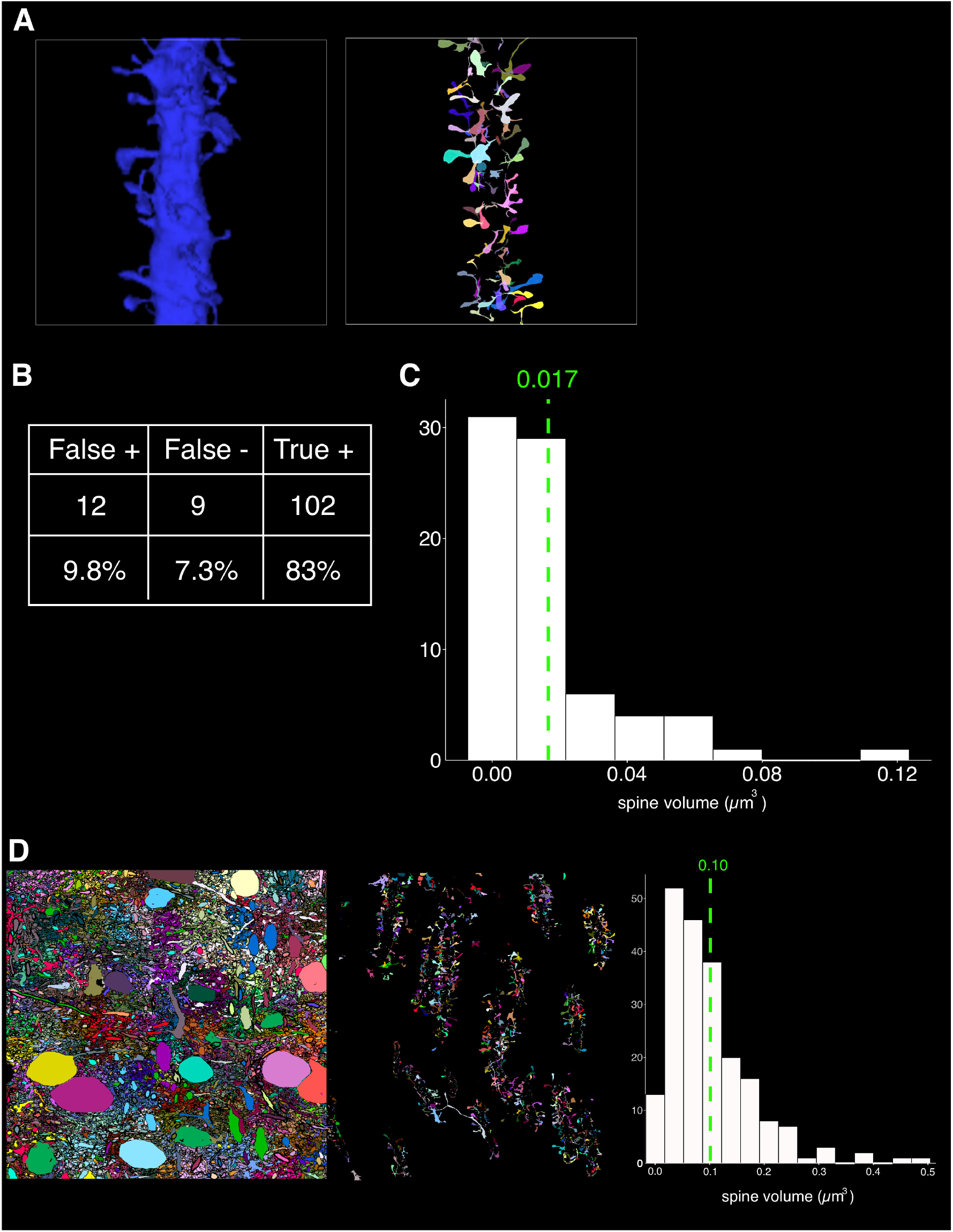
Illustration of the automatic dissociation of spines from spiny dendrites. **(A)** Left panel: Maximum intensity projection of the segmentation of a single spiny dendrite is isolated from the segmented volume. Middle panel: the rendering is converted into a binary image. Right panel: maximum intensity projection of all spines individually isolated and given unique IDs (shown here as unique colors). **(B)** Histogram showing the distribution of spine volumes for a single dendrite in (A); mean spine volume 0.017 ± 0.002 (SEM) µm3, n = 76 spines over 1 dendrite **(C)** False pos/neg **(D)** Left: Single 2D image of all rendered neurons from the full field of view. Middle: maximum intensity projection of all dissociated spines from 10 dendrites extracted from full rendering (left image) and assigned unique IDs. Right: Histogram showing the distribution of spine volumes for 10 dendrites; mean spine volume 0.10 ± 0.006 (SEM) µm3, n = 208 spines over 10 dendrites.

## 4 DISCUSSION

### 4.1 Summary

A key goal in neuroscience is to map the circuitry between populations of neurons at the level of synapses. Presently, the gold standard for studying synapse anatomy is electron microscopy. Recent advances in serial EM offer unprecedented access to studying the 3D anatomy and ultrastructure of synapses along different parts of a neuron (i.e. soma, shaft, spines, etc..), and across numerous cell types. Many technical steps to connectomic analyses from tissue preparation to data analysis are under rapid development by independent research groups, but fast automatic segmentation remains a significant hurdle to making connectomics a readily available technology. As an example, a cubic millimeter of brain tissue imaged at 4nm lateral resolution with 30nm think sections will consist of over two peta voxels and, at current segmentation rates, will consume tens of millions of compute hours. Having fast access to segmentation results will allow users to reconstruct multiple connectomic datasets to understand the role of brain wiring across different comparative axes including development, disease, and evolution. In the current work we aim to overcome this hurdle by scaling publicly available automatic segmentation algorithms. To this end, we show successful scaling of the FFN segmentation algorithm and additionally, we provide an example of how users can leverage the segmentation results by extracting features for morphological analyses (e.g. separating spines from dendrites for spine volume analysis). This demonstration adds to the growing list of computational analysis tools being developed on an open source platform that can be accessed by the greater neuroscience community.

New segmentation algorithms for connectomics are advancing rapidly; however, they often come with increasing demand for more powerful computers, requiring increased processing times and making them less accessible to the general community. Furthermore, as imaging rates increase, producing larger datasets (Yin et al., 2020; Shapson-Coe et al., 2021), this too will put increasing demands for computational resources that can keep pace with these advancements. While advancements in image compression will likely help mitigate the storage load for certain analyses (Minnen et al., 2021), we envision a continuing need for large computing resources as the number and size of datasets grow. We have detailed our solution by leveraging supercomputers at the Argonne Leadership Computing Facility, which utilizes parallelization methods for distributing computationally demanding tasks over hundreds of compute nodes. Specifically, we performed distributed inference using the FFN model on the Theta supercomputer, using both CPU- and GPU-based compute nodes; a future study examining the efficiency of larger-scale supercomputing-based segmentation in more detail is underway. Overall, this pipeline involves fast automation using open source software developed by the connectomics community to relieve the burden of manual annotations combined with a visual interface (i.e. neuroglancer) that allows for scientists to pick out biological features of interest. Importantly, our approach is not tied to a specific algorithm. As new algorithms are developed, we envision a similar ability to ingest and parallelize them to keep current with the latest computational advances being developed by the connectomics research community.

Once data is segmented, we depict a use-case example of how additional image analysis tools can be independently developed using open source tools and easily integrated into our workflow. In our example, we demonstrate a simple method using Python scikit-image tools for isolating spines from spinous dendrites, presumably from excitatory neurons. In doing so, we were able to extract the volumes of spines over an arbitrary number of spiny dendrites. This approach would permit scalable analyses similar to those recently done on spine morphologies (Ofer et al., 2021) but without laborious manual segmentation. As analysis tools become further developed by the broader community (see below), many of these individual features will be automatically parsed and incorporated into meta-analyses similar to commonly used software that parse genome sequencing data into biologically relevant components (i.e. ORFS, ncRNAs, microRNAs, etc.) (Fernández-Suárez and Birney, 2008; Huttenhower and Hofmann, 2010).

### 4.2 Future outlook

To continue advancing connectomics, we anticipate several challenges and opportunities for the field. A major advantage to serial EM is that all cell types and all ultrastructural properties are imaged unlike light microscopy where neurons of interest have to be labeled ahead of time genetically (e.g. GFP) or chemically (e.g. biocytin). Thus, the serial EM data itself is very likely to be useful to the scientific community beyond its original intent. For example, an initial connectomic dataset might be published specifically studying spine density of excitatory neurons, but the same data could be further mined by another research group interested in mitochondrial size distributions. Similar to genomic data, serial EM datasets are thus poised to be a large community resource. To be a useful and reliable resource, however, there are certain considerations that need to be made. Below, we highlight a few, but there are certainly more to follow.

First, accuracy in segmentation is essential, as are metrics for measuring and comparing their success. A variety of segmentation algorithms are being developed in parallel, and we need standardized methods for evaluating which algorithms perform best, and to understand the unique advantages of each algorithm. Having a quantitative method for ranking the performance of different algorithms will ensure that limitations in segmentation are quantified to establish metrics for future improvement. It is noteworthy, however, that even with imperfect segmentation, many biological questions can still be explored such as the spine analysis we outlined above. Second, tools for manipulating segmented data, like those outlined above, need to be advanced and centralized. Centralization can take many forms, ranging from a single website that hosts a catalog of analysis tools to a community-based effort to make independently developed tools publicly available on already established platforms such as Github. Whatever the solution, the connectomics field will benefit from developing such tools with a mindset of open access and user friendliness to ensure data has the maximal impact on the scientific community. Such efforts are already underway and will likely grow as the field develops. Finally, a solution for data sharing, again, ideally in a centralized manner similar to websites such as Neurodata^2^, where the community can access data without having to download and locally store terabyte-sized volumes that often accompany connectomic datasets. For example, building analysis tools from cloud volume segmentations that can be accessed through the web would allow any lab to extract features from a variety of connectomic datasets without having to download and locally store full terabyte (and larger) datasets. These challenges create new opportunities for pushing the field into a high-throughput automated regime that we anticipate will fundamentally shape how we investigate neuroanatomy at the nanoscale.

## AUTHOR CONTRIBUTIONS

Gregg Wildenberg, Tom Uram, and Bobby Kasthuri conceived of the idea. Gregg Wildenberg and Tom Uram designed the figures and wrote the manuscript. Gregg Wildenberg collected the electron microscopy (EM) data which entailed: 1) perfusing the mouse, 2) excising the brain, 3) staining the brain, 4) serial sectioning, 5) imaging, and 6) aligning the data into a 3D image stack. Gregg Wildenberg, Hanyu Li, and Bobby Kasthuri prepared FFN groundtruths. Hanyu Li developed the software packages and performed the distributed segmentation and 3D mesh generation of the full dataset on ThetaGPU. Gregg Wildenberg and Tom Uram performed the multithreaded experiments and segmentation on ThetaKNL. Griffin Badalamente performed the spine separation analysis.

## ACKNOWLEDGMENTS

The authors would like to acknowledge the ALCF Aurora Early Science Program which has supported the computational aspects of this work. Additionally, the authors wish to acknowledge the following members of the Kasthuri lab for their input on the manuscript: Anastasia Sorokina, Vandana Sampathkumar, and Dawn Paukner.

## DATA AVAILABILITY STATEMENT

The datasets for this study are freely available upon request.

## Footnotes

1 https://github.com/google/neuroglancer

2 https://neurodata.io

